# *Mus musculus* mice self-organize existing behaviors into structured bouts as they become expert hunters

**DOI:** 10.64898/2026.09.02.748903

**Authors:** Nishan Shettigar, Ling-Qi Zhang, Jessica Teran, Luisa Schuster, Alex Sohn, Kristin Branson, Emily Jane Dennis

## Abstract

In the wild, a mouse must flexibly perform multiple computations at once, rapidly navigating, sampling its environment, forming and using memories, and balancing internal needs. Capturing the self-paced, low-repetition, complex nature of decision making in the wild while preserving the experimental control needed to interpret its process remains a challenge. Here we show that lab mice, without food restriction, learn to forage for hidden resources cued by ambiguous sounds in a dynamic environment by spatiotemporally reorganizing their existing behavioral repertoire. Using a novel closed-loop assay, we required mice to sit still to receive informative but hard-to-localize sound cues emanating from one correct location they must find, out of 157 possible locations in a large arena. With experience, mice become efficient hunters. Efficient hunting cannot be explained by an increase in a particular behavioral module, but instead, mice reorganize existing behaviors into clustered bouts of high-quality sampling and site-checking. Looking forward, this work establishes an approach for studying the neural, molecular, and evolutionary basis of naturalistic decision-making in mice.

## INTRODUCTION

Like humans, mice are “champion generalists” and live on six continents, thriving in commensal, urban environments like New York City just as well as feral, wild habitats like Skokholm Island^1–3^. Mice live in diverse and dynamic worlds, and must flexibly adjust their behavior to accomplish their multiple, often conflicting goals. These conditions are very different from how we typically study cognition in the lab: training animals to perform hundreds of near-identical trials each hour that isolate a single computation. This reductionist approach has been enormously productive, but it remains unclear whether and how the resulting insights generalize to decisions made in daily life in the wild.

Foraging is a set of behaviors performed in support of the energetic goals of the animal. Hunting is a form of foraging found in predators and omnivores, including *Mus musculus* mice ^2^, a species where we have a wealth of tools available to dissect mechanisms. Searching for prey with ambiguous information is an incredibly rich, self-paced behavior that includes sensation, decision-making, memory, and navigation. As such, hunting provides a rich discovery space for generalizable, interdisciplinary findings. We operationally define hunting as having two phases: “pre-capture”, when mice are searching for a prey item but do not yet know precisely where it is, and this often leads to “prey capture”, which begins the moment mice have a high-confidence estimate of the target’s location and begins to chase, attack, and and kill the prey.

Over the last decade, we have seen a resurgence in studying prey capture in laboratory mice. These studies have provided fresh insights into visual function and active sensation ^4–7^, identification of cell-type-specific circuits and their contributions to behavior^8^, somatosensation ^9^, and motor control ^5^. Together, these studies demonstrate the wide-reaching power of developing a laboratory assay that utilizes a complex, natural behavior. Inspired by this success, we sought to create a self-paced foraging assay focused instead on “pre-capture”, emulating the dynamic, ambiguous, cluttered, and rich environments in the wild.

## RESULTS

### Mice learn to forage for hidden prey in sound-guided foraging task using Modulo: a flexible, closed-loop behavior arena

Towards this goal, we built a large (2.29 x 1.97m) hexagonal arena made of 157 modular hexagonal tiles and 24 half tiles at the edges (Figure 1A). We use two hexagonal tile types: plain tiles that are empty inside of the felt walls, or release tiles that contain a speaker and a trap door (Figure 1B). The trap door can be opened by a pneumatic system controlled by the main computer, and usually contains a tethered cricket as a reward (Figure 1C). The tether is tied to a post, and ensures that after we open the trap door, the cricket cannot escape the mouse nor the tile to which it was assigned. The speaker plays sounds on command, driven by a Teensy controlled by the main computer. Regardless of type, each tile contains six posts which are wrapped in a loop of felt to create walls that obfuscate the interior of the tile from the point of view of a mouse running within the corridors formed by the walls of adjacent tiles (Figure 1D). Experiments were run under low-light conditions (~4 lumens), providing limited access to visual cues (Figure 1D).

**Figure 1:**
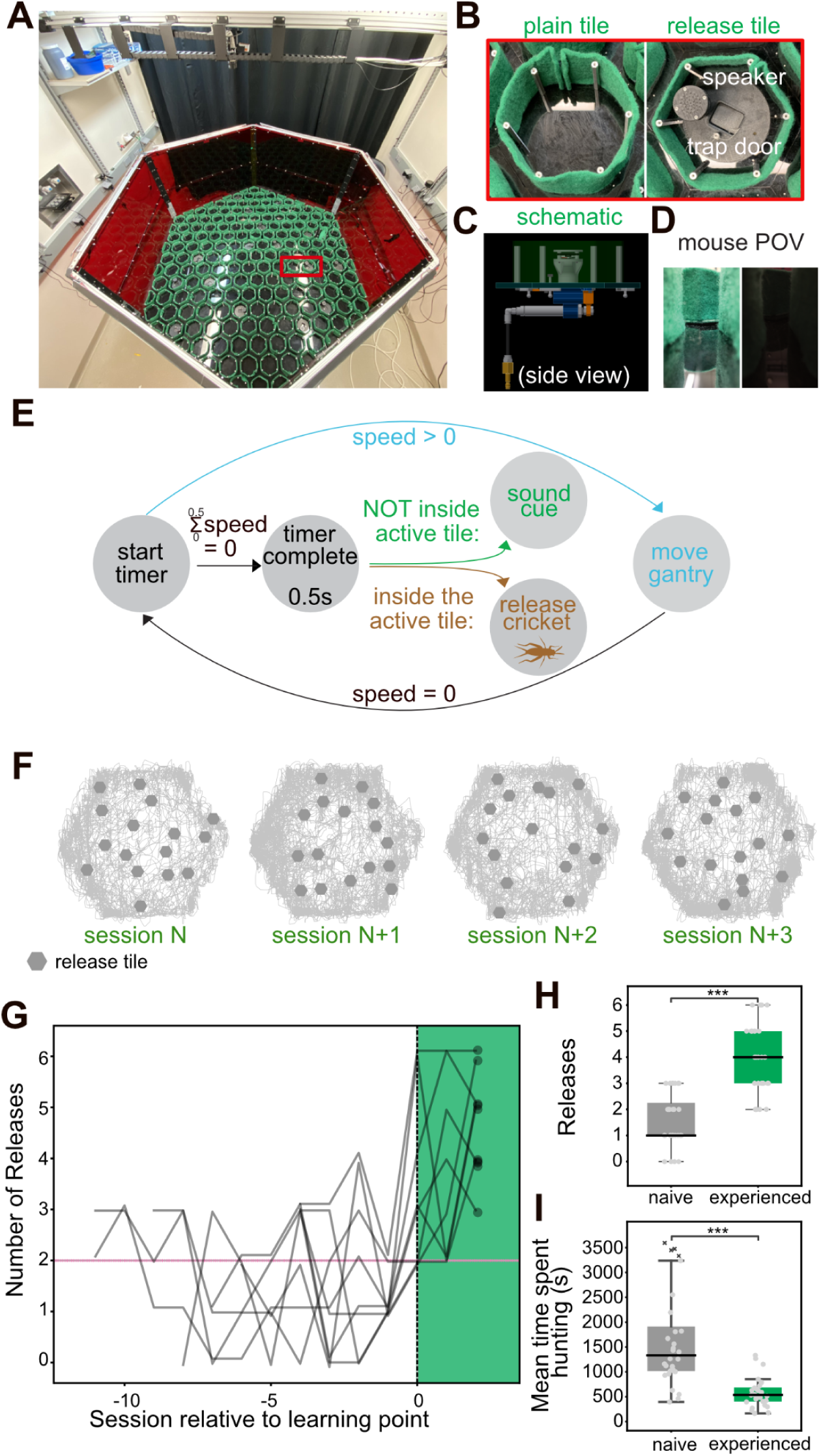
Mice learn to forage for hidden prey in a sound-guided foraging task using Modulo: a flexible, closed-loop behavior arena. (A) Image of Modulo, a large, reconfigurable arena consisting of a motorized, overhead, three-axis gantry in which two axes are linked in lockstep to form a two-dimensional gantry system. In real time, tracking camera data drives the two axes to adjust the camera positions, so they remain above the mouse. The hexagonal arena comprises 157 hexagonal tiles with green wool felt walls. Red box highlights tile types shown in detail in B. (B) Images of the two tile types: plain tiles (left) and release tiles (right). Each tile is bordered by a felt wall, and the spaces between adjacent tiles form the corridors of the maze-like arena. Release tiles contain a trap door and a speaker, with sounds driven by the closed-loop system. (C) Schematic of a release tile. Beneath the trap door, release tiles contain a concealed sub-floor chamber, enabling the release of prey (crickets, white) or other items. The trap door is pneumatically controlled. (D) Images from the point of view of a mouse in Modulo’s corridors, well-lit for demonstration (left) or under our experimental conditions (right). (E) Schematic of the closed-loop workflow during hunting sessions: real-time tracking of the animal allows the animal to control sound cues and trigger cricket release by opening the trap-door when the mouse is in the correct location. (F) Representative movement trajectories from a single animal across four hunting sessions. Note that the sixteen release tiles (grey) change location between sessions. (G) Mice learn the task in 5–12 sessions (median = 8.5). We consider mice experienced after two or more releases per session for 3 successive sessions, with at least one four release session. We aligned the release data for each mouse to its experienced point, and the shaded band marks experienced sessions. (H) Number of releases per session compared between naive (first four sessions per animal) and experienced sessions (8 animals; n = 28 sessions per group; two-sided Mann–Whitney U test, p < 0.0001). (I) Mean time spent hunting per release compared between naive and experienced hunters; × indicates sessions with no release (8 animals; n = 28 sessions per group; two-sided Mann–Whitney U test, p < 0.0001)

After acclimating mice to human handling^10^, crickets, and the arena, mice are provided the full task without any shaping. At any given moment, there is only one goal location out of 16 possible release sites. The mouse must find the correct goal location, cued by sounds that play from the speaker at the goal location. Mice control when and where they receive this information: if a mouse sits still for at least half a second, we play a sound from the speaker at the goal location. No sounds play when the mouse is moving. If the mouse enters the goal location and sits inside the walls for at least 0.5s, we open the trap door and release the cricket (Figure 1E, see also Supplementary Figure 1). All release tiles are moved between sessions, so mice cannot use the locations of release sites from day to day (Figure 1F).

Mice often release at least one cricket in their first hunting session, but improve with experience (Figure 1G). With days of experience, mice increase the number of crickets (Figure 1H) and become more efficient: they are able to find more crickets per session and spend less time finding each cricket (Figure 1I). Both male and female mice improve with experience, but we focus our results on female mice, because they learn more quickly (Supplementary Figure 2). Further, we were unable to consistently co-house all males, and did not want to study the effects of social isolation stress^11^.

### Mice require sound cues to efficiently find hidden prey

The sounds playing from the speaker are the only *unique* cue indicating the correct goal location. However, mice could solve the task without using the sounds by finding and checking all of the release tiles. In familiar environments, mice explore novel objects and spend more time exploring objects that can be climbed, like our speakers^12^. These proclivities likely explain why, in our task, even naive mice visit cricket release tiles (Figure 2B). We define “tile checks” as events where an animal enters a release tile, and waits at least 0.5 seconds: this is long enough to trigger the trap door if they are in the goal location. Mice slightly but significantly increase the number of “tile checks” as they gain experience (Figure 2B). This is an indication that they learn the association between the trap door opening and waiting inside the walls of a release tile. However, experienced mice collect rewards with far fewer “tile checks” than expected by chance, therefore they must be using other cues in addition to tile location (Figure 2C). We then tested which cues were required for experienced mice to find and release crickets. We measured the probability of mice finding the goal location, the time it took for mice to find the goal location, and the distance travelled to successfully release the cricket. Removing the cricket from the goal entirely and decreasing visual cues by removing all visible light sources had no significant effect on any of these metrics. Food-depriving the mice also did not have a significant effect on these measurements of success. In contrast, when we do not play any sounds, which are the only unique cue indicating the correct goal location, mice find the goal less often (Figure 2D), and when they are able to find the goal it takes them longer (Figure 2E) and they must travel a longer distance (Figure 2F). Sound cues are therefore necessary for experienced mice to efficiently find rewards.

**Figure 2:**
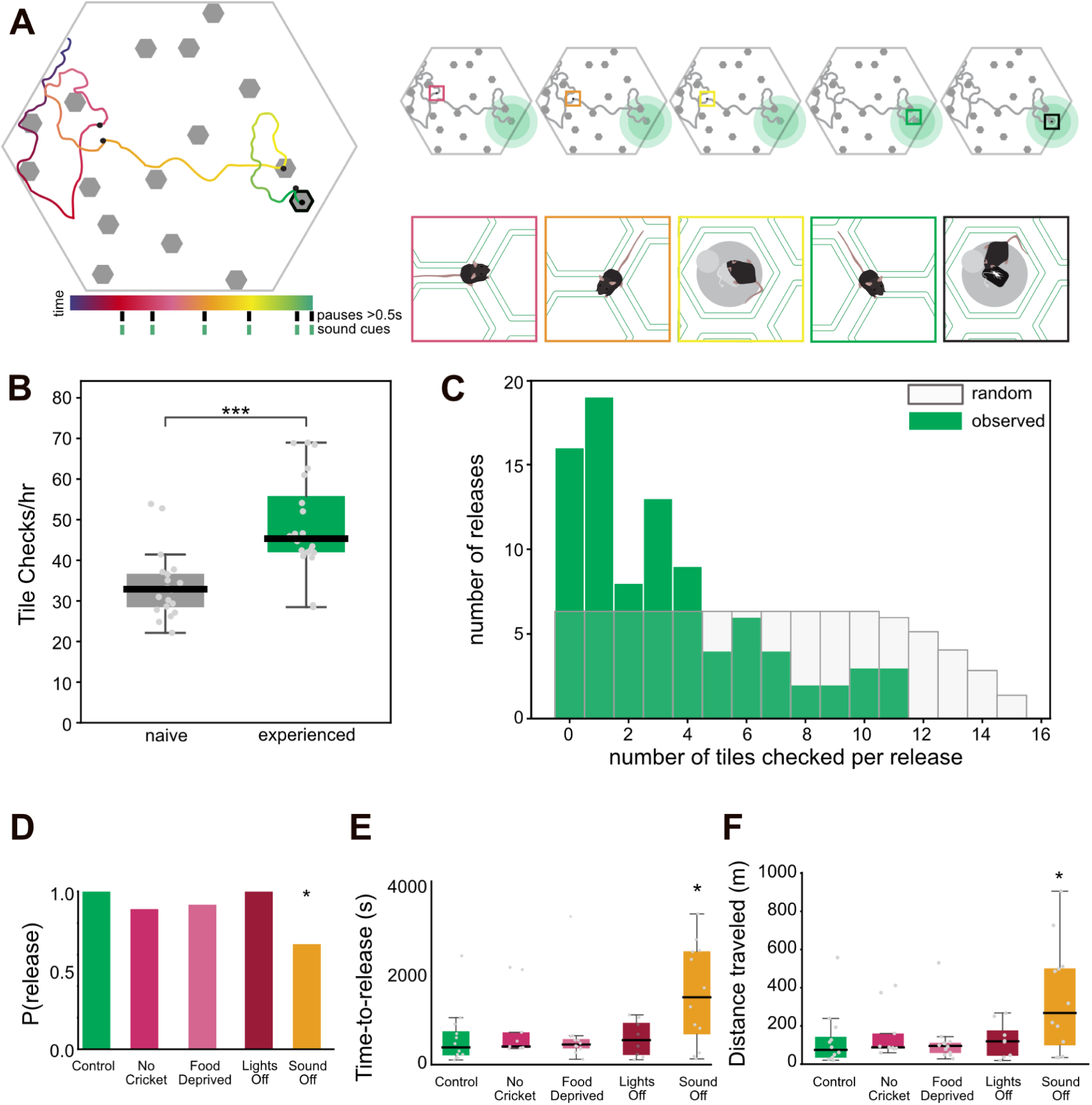
Mice require sound cues to efficiently find hidden prey. (A) Schematic of the task, and a hypothetical trace highlighting a tile check (yellow), several pauses (pink, orange, green), and a final pause in the correct location, resulting in a successful cricket release (black). (B) Number of tile checks (animals entering a release tile and receiving a sound) compared between naive and experienced sessions. Boxes show the median and interquartile range (IQR) with 1.5×IQR whiskers; grey points are individual sessions. Tile checks increased significantly with experience (6 animals; n = 20 sessions per group; two-sided Mann–Whitney U test, p < 0.0001). C) Distribution of wrong-tile checks before trigger per release in experienced animals (purple; 89 releases from 6 animals), shown against the distribution expected under random search (grey). Random search was modeled as sampling distinct release tiles in uniformly random order from the remaining pool until the goal tile; for a release with P available tiles, the number of wrong-tile checks is uniformly distributed over 0 to P−1, and the grey bars show the resulting expected counts summed across all releases (computed analytically). Significance was assessed on the mean wrong-tile check count by a one-sided per-release permutation test, drawing one random count per release from its own pool size and repeating 10,000 times (p = 0.0001). (D) Without sound, the probability of a mouse releasing the cricket at second location decreased relative to all other conditions (one-sided Fisher’s exact test, p = 0.046). (E) Time to second release across conditions. Boxes indicate the IQR, whiskers 1.5×IQR, the black line the median, and grey points individual sessions. Sound Off increased the time to second release relative to control (two-sided Mann–Whitney U, p = 0.040). Non-releases were scored as time to session end. (F) Distance travelled (m) to the second release across conditions. Boxes indicate the IQR, whiskers 1.5×IQR, the black line the median, and grey points individual sessions. Sound Off increased the distance travelled relative to Control (two-sided Mann–Whitney U, p = 0.046). Non-releases were scored as distance travelled to session end. For (D)–(F): 4 animals, n = 12 Control, n = 9 No Crickets, n = 12 Hungry, n = 10 Lights Off, n = 12 Sound Off.

### Efficient foraging is not explained by an increase in the frequency of behavioral modules

Sounds are required for mice to hunt efficiently, and mice have control over when and where they trigger sounds from the goal location. We asked if mice increase the frequency of sound-gathering behaviors as they become more efficient hunters, but instead found that mice do not significantly increase the frequency of sounds triggered per session (Figure 3A). However, in this self-paced paradigm, mice are able to engage and disengage from the task at any time during the session, and may be triggering sounds when they are not actively hunting. Further, mice can trigger sounds with any behavior that keeps its centroid still for at least a half of a second, and this can include task-irrelevant behaviors like grooming and task-relevant but not searching-relevant moments, like when the mouse is eating the cricket (Figure 3C). To identify these different sound-triggering behavior, we used our high-speed (120 fps) camera data to track 37 keypoints on the mouse’s body. These keypoints were used to classify sound-triggering behavior into different pose categories (Figure 3B) using a supervised machine learning procedure (APT and JAABA^13,14^).

**Figure 3:**
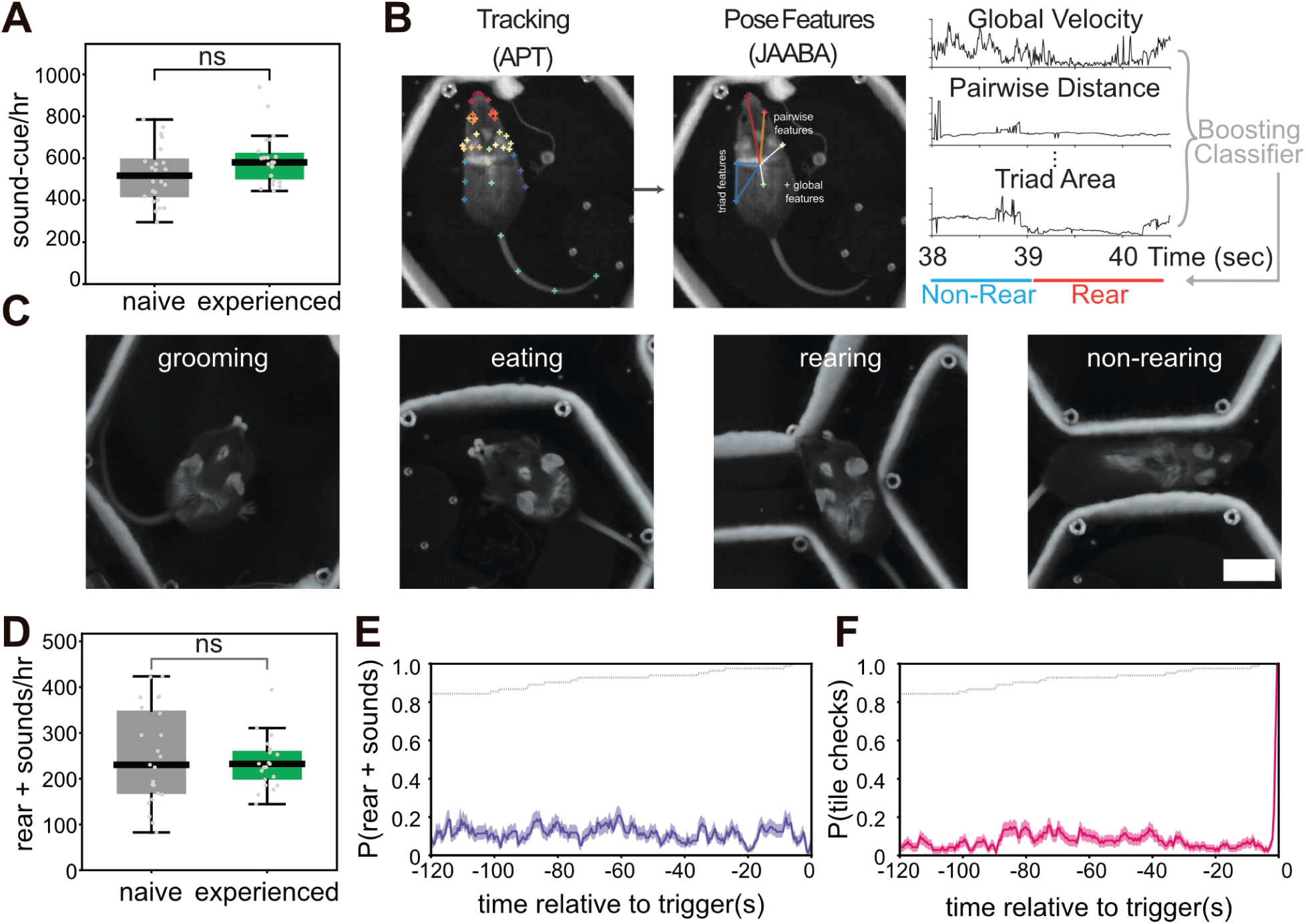
Efficient foraging is not explained by an increase in frequency of behavioral modules. (A) Sound-cues per hour compared between naive and experienced sessions; no significant difference. Boxes indicate the IQR, whiskers 1.5×IQR, the black line indicates the median, and grey points individual sessions (6 animals; n = 20 sessions; two-sided Mann–Whitney U test). (B) Pipeline for automated detection of rearing behavior. Body landmarks are tracked from high-speed video using APT (left). From the tracked keypoints, JAABA-style features are computed (middle) - pairwise features (distances between landmark pairs), triad features (areas defined by landmark triplets), and global features (e.g., overall velocity). The resulting time series of features (right; Global Velocity, Pairwise Distance, Triad Area) are fed to a boosting classifier that labels each frame as Rear or Non-Rear (colored bar, bottom). (C) Representative high-speed camera frames of animal poses during sound cues, illustrating the behavioral categories scored (left to right: grooming, eating, rear, non-rear). White scale bar = 2 cm. (D) Rear+sounds per hour compared between naive and experienced sessions; no significant difference. Boxes indicate the IQR, whiskers 1.5xIQR, the black line indicates the median, and grey points are individual sessions (6 animals;naive, n = 23 sessions; experienced, n = 21 sessions; two-sided Mann–Whitney U test) (E) Peri-release probability of rear+sound events. For each release, the probability of the animal being engaged in a rear+sound is plotted as a function of time relative to the release (release trigger at 0 s; window extending to 120 s before release). The probability is availability-normalized: at each time point it is computed over the hunts with data at that time (proportion of total hunts indicated by grey dotted line). N = 83 releases from 6 animals. (F) Peri-catch probability of tile checks (release-tile visits containing a sound cue), computed and plotted as in (E) over the same 83 releases from 6 animals. In B and C, images were cropped and adjusted to increase brightness 40% and contrast 100% to improve visibility.

In particular, we set out to create a classifier trained to target grooming. From above, eating and grooming recruit similar movements, and we allowed our classifier to identify both grooming and eating events, as both grooming and eating are unlikely to be a part of active search. Anecdotally, we also noticed that our experienced animals rear often (Figure 3C). Rearing is an exploratory, investigative posture that mice often deploy when gathering information^15,16^. We trained a separate classifier to identify these rearing events. We used these data to classify each sound cue as ‘rear+sound’ or ‘sound-cue’. Like sound-gathering, we find that rear+sound events do not increase with experience (Figure 3D).

Therefore, sounds are required for efficient performance, but the frequency of sound-gathering behaviors does not increase with experience. Similarly, tile check frequency increases, but this alone cannot explain the efficiency of our mice. Because our task is self-paced, it is possible that mice are only actively hunting for short, focused periods during each session, and therefore these session-wide metrics may be insufficient. We reasoned that experienced animals should be engaged in the task just before successfully releasing a cricket at the goal location, and therefore focused our analyses only on the two minutes before each successful cricket release. Surprisingly, we found that there is still no significant increase in the frequency of sound-gathering, rearing during sounds, nor tile checking in the two minutes before cricket releases (Figure 3E-F), despite the sounds and tile checks being important components of successful hunts.

### Efficient foraging emerges from animals restructuring their existing behavior modules into bouts

Perplexed, we re-visited our assumptions. In highly trained tasks, animals must typically learn an association between a stimulus and a behavior they would otherwise rarely express: licking, poking, or lever-pressing. Mice in these tasks greatly enrich these behaviors with learning. However, even naive mice in our arena already readily express all of the behaviors needed for success: exploring, rearing, pausing, and entering tiles. To find the hidden crickets, mice do not necessarily need to do these behaviors more often, they just need to deploy them in a targeted way. We therefore looked for time periods where mice transitioned between rearing and tile checking. We defined “engaged” bouts where mice chain together rears and tile checks in quick succession (Figure 4A).

**Figure 4.**
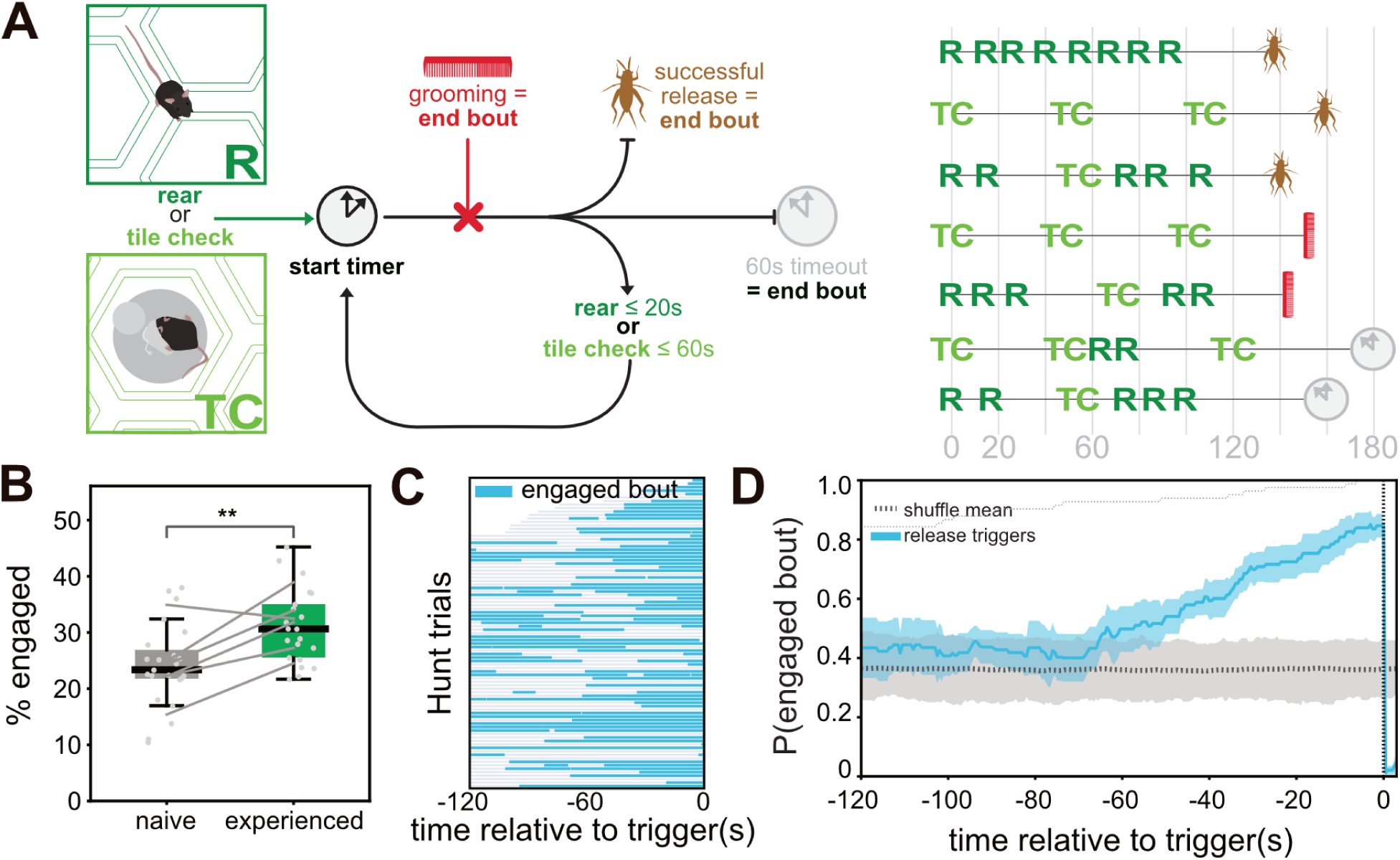
Efficient foraging emerges from animals restructuring existing behavioral modules into bouts. **(A)** Definition of an engaged bout. Left: schematic of the bout-construction logic. Engaged bouts are contiguous chains of rear (R) and tile-check (TC) events, poses during which the animal received a sound cue, linked by a timer-based rule and terminated by grooming or eating, a successful release (capture), or a timeout (see Methods). Right: example event sequences of single bouts, aligned to onset (time in seconds), each ending in a capture (cricket), a grooming/eating episode (red comb), or a timeout (gray clock). **(B)** Percentage of session time spent in engaged bouts, compared between naive and experienced animals. Each point is one session; boxes indicate the IQR, whiskers 1.5×IQR, and the black line the median. Lines join the per-animal median naive and experienced values (6 animals). Time in engaged bouts was higher in experienced animals (6 animals; n = 23 naive sessions; n = 21 experienced sessions; two-sided Mann–Whitney U test, p = 0.0036) **(C)** Raster plot of engaged bouts aligned to release triggers, showing 83 releases from 6 animals across 21 sessions (time axis extending to 120 s before release). Each row is one hunt, ordered by duration; blue segments mark engaged bouts. (D) Peri-release probability of engaged bouts as a function of time relative to the release trigger (release at 0s; window extending to 120s before release), across 83 releases. The probability is availability-normalized: at each time point it is computed over the hunts with data at that time. The shuffle control (grey) is overlaid: release times were redrawn uniformly within each session (count preserved, bouts unchanged); the dotted line indicates the shuffle mean and the band the 2.5–97.5th percentile of shuffles.

Using this approach, we found that mice increase their engaged bouts with experience (Figure 4B), engagement is enriched just before successful releases (Figure 4C), and animals are significantly more likely to be in an engaged hunting bout for the 60s prior to a successful cricket release (Figure 4D). Engaged hunts explain over 85% of all observed releases, and the definitions and results are robust to the parameters used in the definitions (Supplementary Figure 3).

## DISCUSSION

In this study, we demonstrate that mice, without food restriction, learn to locate hidden food rewards signaled by ambiguous sound cues in a changing environment by spatiotemporally re-organizing their behavioral repertoire. Although this study is entirely behavioral, our experiments are amenable to electrophysiology, imaging, and perturbations: all of our experiments are by default performed in closed-loop, and orchestrated by ONIX hardware^17^ and Bonsai software^18^. These tools were designed to facilitate freely moving experiments: mounting a commutator to the 2D gantry allows us to perform tethered recordings and perturbations in a large arena without the drawbacks of a fixed tether in a large space and without the downsides of using the heavy batteries required for long recordings using wireless designs. By using a moving gantry system, the commutator is always directly above the mouse’s head regardless of its location, and therefore there will be no tension from being far away from the tether. In addition to this design feature, below, we draw attention to additional key features of our work that extend its relevance beyond our immediate findings, opening avenues for investigation across multiple subfields of neuroscience.

### Active sensation

In our study mice physically sample the environment and use sound cues to find hidden locations containing food rewards. Active sampling has been studied in rodent olfaction as sniffing^19,20^, vision through eye and head movements^21,22^, and somatosensation through whisking^23^ for decades. A resurgence in studying active sampling in rodents through ethological behaviors has already revealed new insights into neural encoding of stimuli^24^, feature extraction^8,25^, and has revealed that active sensation shapes how sensory information is used to support other behaviors, decisions, and computations ^26^. Hearing is also an active process ^27^, yet there are comparatively few paradigms to study active sensation in the auditory system alone. To address this gap, Mai and colleagues^28^ developed a sound-seeking task to examine how movement enhances perception and how animals recover from sensory loss and disorders. As in our task, mice also adjusted their posture and exploration to sample the sound cues. A common thread in both experiments is that mice can exploit the physical structure of the environment to simplify the task, harnessing the behavioral flexibility freely living animals can exhibit to solve problems in the wild. However, our work is distinct in that our target locations are dynamic and change daily, and mice must find hidden targets over large distances. These differences encourage different physical solutions, and it will be interesting to compare results across these two tasks in the future to better understand how mice encode and use ambiguous sound cues in different complex environments.

### Self-paced, self-organized, and self-motivated

Our task is also self-paced and self-organized: mice learn the task without any explicit shaping or training, can engage or disengage in the task at will, and animals are able to organize their movements and behaviors however they choose. Because of this, no two trials are identical. This was a part of our behavioral design, as we wanted to capture the diversity of movements and behavioral flexibility of wild mouse behavior as mice pursue a goal. This exciting behavioral aspect may pose a challenge when considering neural activity in the future. However, a recent study gives us cause for optimism: here, researchers played hide-and-seek with rats while recording from dozens of neurons in the rat’s prefrontal cortex ^29^. In this study and in a follow-up study, they found neural correlates of task variables at both the individual neural level and at the population level from a relatively small number of cells ^30^. Similarly, in freely moving, foraging macaques, identifying meaningful moments was sufficient to discover neural correlates of behavior^31^ and coding of strategic variables like reward and choice ^32^ by focusing analyses on moments where the macaques were engaged in the task.

Notably, in our task, animals do not need to be food-deprived nor water-deprived. In the wild, and even in homecages in the lab ^33^, mice do not wait until they’re starved to search for nor consume food. In the visual system, food and water restriction can influence task engagement and performance in a discrimination task ^34^, and caloric restriction decreases the coding precision of visual stimuli ^35^. In future work, we can build on these findings from the visual system and compare sated mice, calorically restricted mice, and acutely food-deprived mice behaviorally and neurally to explore how energetic limitations can affect neural encoding in the auditory system during behavior.

Here we demonstrate that mice can learn to forage for hidden, acoustically-cued resources in a dynamic environment, not by acquiring new behaviors but by spatiotemporally restructuring their existing behavioral repertoire. Together, our results provide a platform for studying the behavioral, neural, and molecular basis for self-paced, sensory-guided decision making, and an example of how natural behavior can continue to surprise us.

## AUTHOR CONTRIBUTIONS

Conceptualization, N.S., J.T., E.J.D.; Data Curation, N.S., L-Q.Z., J.T., E.J.D.; Formal Analysis, N.S., L-Q.Z., K.B., E.J.D.; Investigation, N.S., J.T., L.S., E.J.D.; Funding Acquisition, E.J.D.; Methodology, N.S., J.T., A.S., E.J.D.; Software, N.S., L-Q.Z., K.B., E.J.D.; Visualization, N.S., L-Q.Z.; Writing – original draft, N.S., L-Q.Z., E.J.D.; Writing – review & editing, N.S., L-Q.Z., J.T., L.S., K.B., E.J.D.

## ACKNOWLEDGEMENTS

We thank Kerry Sobieski for project administration; Lakshmi Narayan for technical assistance setting up the Zaber system; Frank Loesche, Carmon Morrow, and the Mechanistic Cognitive Neuroscience NeuroEngineering Team for assisting with system synchronization; and John Macklin and Janelia Experimental Technologies for additional technical assistance. We also thank Ahmed El Hady, Alison Comrie, Ben Arthur, Jakob Voigts, Roian Egnor, and Samantha Moseley for project feedback. This work was supported by the Howard Hughes Medical Institute, where EJD was a Group Leader at the Janelia Research Campus.

## METHODS

### Experimental model and study participant details

All work was performed at Janelia Research Campus in Ashburn, VA under IACUC protocol 24-0263 “Neuronal investigations of cricket hunting in *Mus* mice” and conform to all applicable regulatory standards.

#### Mice

We used 8 female and 2 male wild-type *Mus musculus* mice ages 2-18 months. Four were B6.CAST-*Cdh23^Ahl^*^+^/Kjn (RRID:IMSR_JAX:002756) and 4 females and 2 males were PWK/PhJ (RRID:IMSR_JAX:003715). All animals were kept on a 12h:12h inverted dark:light cycle, and all experiments were run during the dark phase for the animals, and animals run between hour 13-19 in the dark phase. Mice were provided mouse chow *ad libitum* in their homecages unless otherwise described (LabDiet 5053) and cages included corncob bedding and enrichment items including at minimum nesting material (Bed-R-Nest or nestlets), a place to hide (plastic tunnel or running wheel with dome). All cohorts of animals run on the same day were animals from the same cage except males, who originated from the same cage. We bleached a T pattern and a dot on the head of our mice approximately every other week (Manic Panic Flash Lightening Hair Bleach) to assist live tracking, and each mouse received an RFID chip to assist in identification (UC-1283 MiniMax Programmable Microchip, 1.2×8.3mm). After any exposure to anesthesia, mice were given at minimum 24 hours to recover before performing any behavioral experiments. Mice were housed in a free-standing ventilated rack (Allentown Inc) ventilated with 100% outside filtered air.

#### Crickets

All crickets used were female or juvenile *Acheta domesticus* (Fluker Farms) or *Gryllodes sigillatus* (Ghann’s Crickets, Augusta GA #C14 #C38) and were ¼-1½ inch in size. In these species, only adult males can chirp because females and juveniles do not have wings required to make the noises. All crickets were provided with bedding made from a 75/25 mix of 75% Fluker’s High-Calcium Cricket Diet (Fluker Farms) and Organic Vermiculite (Fluker Farms) and fed Fluker’s Cricket Quencher (Fluker Farms). We also provided paper egg filler flats for hiding (ULINE S-5189) in an acrylic plastic tank. Crickets were tied with food safe butcher’s twine.

### Behavior

#### Acclimation to humans and crickets

Upon arrival, animals were allowed to acclimate to the housing room at least 24 hours. Animals were then subjected to a 5-10 day acclimation protocol ^10^, first introducing them to human presence (1-2 days), physical interaction (2-5 days), and handling (2-4 days) until comfortable with human interactions, defined by eating and exhibiting relaxed postures when handled.

Next, mice were acclimated to crickets. We removed all cage mates and items from their homecage and introduced a cricket for 5 minutes. Each mouse received at least 3 individual crickets. We continued exposing the animals to crickets until they chased, captured, and killed a cricket in their home cage.

#### Modulo

##### Tiles

All tiles were made of cast acrylic (McMaster 8505K742), laser cut to size (15.1×17.4cm hexagon), and affixed with push-to-open grab latches on the bottom (McMaster-Carr 10825A28) and 1.75” threaded hex standoffs on top (McMaster-Carr 91780A632). The “walls” of each tile were made out of a loop of 100% wool felt (1/10in thick, Spring Valley, Weir Crafts National Non-woven) sewn together with organic cotton thread (Gutermann).

Cricket release tiles additionally have a 3D printed insert (FormLabs Black Resin) with a 3D printed casing for a speaker (PUI Audio AS02704MS-N50-LW100-R) and a 3D printed trap door. The speaker is connected via audio jack to an Audio Shield (Rev D) and Teensy 4.0 receiving commands through a microUSB connected to the main computer. The trap door was controlled by signals sent to a separate microcontroller (PoRelay8 v1.1) sending voltage signals to a pneumatic valve coupled to the trap door through polyurethane tubing and a push-to-connect adapter (McMaster-Carr 5779K104, 5648K23).

##### Gantry and payload

We assembled a two-dimensional tracking system made of three motorized moving stages (2.25m LC40 with dual-motor axis, Zaber Technologies, see Supplementary Figure 1). Two cameras (Basler ace acA2040-180km NIR) each with an Edmund optics lens (25mm, 50mm) were mounted to one of the axes. One streamed to the main computer to be used by Bonsai for live animal tracking. The second camera was connected to a frame grabber (microEnable 5 marathon ACL FrameGrabber, Basler) on a separate computer and streamed 120fps, 1024×1024 pixel video directly to disk on a separate computer for post-processing. The frame grabber emits a TTL pulse through a customized BNC cable connected to the ONIX system ^17^ allowing for synchronization across computers and devices. A torque-free commutator (OEPS-7759) was also added to the gantry payload alongside the cameras, though not used in this study.

##### Bonsai

All aspects of the experiment were controlled through Bonsai ^18^ and external devices were synchronized either through USB or the ONIX PCIe Acquisition System. See Supplementary Figure 1 for details. To track the animal in real time, we used SLEAP 1.2.9 ^36^ or DeepLabCut-Live! ^37^ with a frame rate of approximately 15fps. The animal’s nose and tailbase, and 5 bleached points on the animal (dot on the head, and a T on the back including the top left of the T, where the T crosses itself, top left of the T, and the base of the T) were tracked and a median of at least two high-quality points was used as the animal centroid. We calculated the euclidean distance between the center of the image and the mouse’s centroid, and used this value to drive the gantry system towards the mouse’s location. Pauses (stillness) were calculated as the mouse’s centroid needed to be less than 1/14 - 1/16 from the center of the image, adjusted manually based on the mouse’s size. To trigger either the trap door to open (cricket release) or a 72+/-2 dB SPL sound (0.4 volume) to play from the speaker in the goal location, the centroid of the animal must be tracked and not leave a radius of 1/14-1/16 of the center of the image, for at least 0.5s. To open the trap door, the animal must further have been within 7200 Zaber units (10.9cm) of the center of the correct release tile location.

##### Environmental measurements

To measure speaker variability, we used a laptop with Arduino’s (version 1.6.1) to deliver commands. For a given tile, the tile’s speaker was connected to the teensy audio shield, and an Extech 407730 digital sound level meter was placed at a 90 degree angle from the speaker shield surface, with the windshield barely touching the speaker shield. All tests were done in a 54dB room. Each software volume value (0.1, 0.2, 0.3, 0.4, 0.5, 0.6, 0.7) was played 3 times in succession. We used the ‘A’ frequency weighting, and ‘FAST’ measurement (125ms measurements). See Supplementary Figure 4. We chose a software volume of 0.4, corresponding to 72+/-dB SPL and checked the release tiles for volume accuracy.

Using a Vibration meter and stethoscope (McMaster #8738T1, vibration range of 10-10000Hz) we were unable to measure any velocities (0.01-19.99 in/sec) nor accelerations (0.01-19.99g) except on the speaker itself (0.02 in/sec). We measured vibration at the speaker shield, the 3d printed insert (which holds the trap door and speaker housing), and the acrylic of the cricket release tile while playing sound cues. Mice are sensitive to 10-200 Hz vibrations above 0.05 m/s^2^ ^38^ and deafened mice show no behavioral response to 500-10000Hz sound played between 60-80 dB SPL in a swim task, but are sensitive to 6-50Hz when played at 160-175dB ^39^. The sounds we produce start at 3000kHz and below 80dB so we did not expect vibrations to play an important role in this task, but took these measurements to ensure our expectations were correct.

To measure the luminance in the arena under different conditions, we used light meter 14855K649, McMaster Light meter for White LEDs and found the arena was 3-5 lumens at all locations when white lights were on at 8V, our most common experimental condition. The light meter could not measure any lumens when our white lights were off (0V, Lights Off condition).

##### Modulo Experiments

After acclimating to humans and crickets, mice were introduced to the Modulo arena. Each animal experienced at least four days of acclimation, each day completing a single 40-65 minute session. During this time, no sounds were played and no cricket release tiles were provided.

After acclimation, animals were provided with the full task. Each day 16 cricket release tiles, each containing a live, tethered cricket, were placed in semi-random locations. We regularly evaluated previously used tile locations and ensured approximately even coverage of the arena. Mouse order was shuffled frequently.

##### Modulo Manipulations

In the No Cricket, Food Deprivation, and Lights Off manipulations, the experimental logic remained identical to the control task. In the No Cricket manipulations, no cricket was added to the second correct release location. For Food Deprivation experiments, all chow was removed from the home cage 16–24 hours prior to behavioral testing. For Lights Off experiments, all light sources were turned off for the entire session. For Sound Off experiments, the speaker at the second reward location was disconnected from the Teensy at the start of the session, so that no sounds were produced there while the animal searched for the second reward after successfully triggering the first.

To quantify the effect of each manipulation on the search for the second reward, we measured the time taken and the distance travelled to the second release. For each session, the analysis window opened 30 seconds (the lockout interval) after the first release and closed at the second release. Distance was computed as the cumulative tracked path length over this window. In sessions where the animal never reached the correct second reward location, the window was closed at the end of the session, so that the full time spent and distance travelled while searching for the second reward were included in the analysis.

## QUANTIFICATION AND STATISTICAL ANALYSIS

All quantification and statistical analysis were done in Python [3.8.18]^40^, using the following modules: NumPy^41^, Pandas^42^, SciPy^41^, Statsmodels^43^, Seaborn^44^, and Matplotlib^45^. Specific details of the statistical test, number of samples, definition of n, and p-values are given in the figure legends. Unless otherwise stated, box plots show the median (center line), interquartile range (IQR; box) and 1.5×IQR whiskers, with individual sessions overlaid; peri-release curves show the mean ± SEM.

Session-level comparisons between naive and experienced animals used the two-sided Mann–Whitney U test. The probability of a release was compared using a one-sided Fisher’s exact test with the direction pre-specified. For the manipulation experiments, the Sound Off condition was a pre-specified primary hypothesis and was compared against control using the two-sided Mann–Whitney U test, reported without correction. The other conditions were likewise compared against Control by Mann–Whitney U test with Holm–Bonferroni correction; none differed significantly. Significance in figures is indicated by asterisks (*p < 0.05, **p < 0.01, ***p < 0.001); non-significant comparisons are labeled “ns.”

Observed wrong-tile checks were compared against a random-search baseline. Random search was modeled as sampling distinct release tiles in uniformly random order from the remaining pool until the goal tile, so that for a release with P available tiles the number of wrong-tile checks is uniformly distributed over 0 to P−1; the expected distribution was computed analytically across all releases. The observed mean wrong-tile count was compared against this baseline using a one-sided per-release permutation test, drawing one random count per release from its own pool size and repeating for 10,000 iterations.

No statistical test was used to predetermine necessary sample sizes. Sessions were rarely excluded, and all exclusions were due to technical issues (e.g. the release tile would not open when triggered) or if the session did not have all data streams for at least 40 minutes.

### Keypoint tracking (APT)

We conducted high-resolution keypoint tracking of the animal through the high-speed camera, recorded at 120 frames with 1024 x 1024 px resolution. Thirty-seven landmarks (nose, eyes, ears, back, tail, and body axis, see Supplementary Figure 5) were manually labeled in Animal Part Tracker (APT) ^13^ on full resolution grayscale frames. We labeled a total of 799 frames of mice across all strains and sexes used in our experiment across a variety of posture conditions, drawn from 253 video clips.

The landmarks were tracked with DeepLabCut ^46^, implemented with a ResNet-50 backbone initialized from ImageNet-pretrained weights, plus the DeepLabCut location-refinement head. Training data were augmented on-line using APT’s augmentation pipeline, which applies random rotation, translation, scaling; and brightness and contrast jitters. The network was trained with stochastic gradient descent with momentum (0.9), batch size 16, weight decay 10^-4^, following a multi-step learning-rate schedule (0.005 for the first 10,000 iterations, then 0.02 to 430,000 steps, and 0.002 afterwards). The model used for all analyses was trained a total of 732,000 iterations. Training was performed on the Janelia compute cluster and took approximately 12 hours on a single NVIDIA A100 GPU.

To evaluate the performance of our tracker, we performed a five-fold cross-validation procedure. In each fold, 80% of the frames were used as training data and the remaining 20% as test data, with the model architecture and training procedure exactly as described above. This was repeated five times so that each frame served as a test frame exactly once. For each hold-out frame, we computed the error as the Euclidean distance between the predicted keypoint and the corresponding human label. The resulting cross-validation errors are visualized as follows: Tracking performance was accurate for landmarks on the nose, eyes, ears, neck and back, with median errors all below 5% of body length. Tracking was less reliable for the tail section and the sides of the body (median errors of 12.1% and 12.5% of body length, respectively). (Supplementary Figure 6)

### Behavior Classification (JAABA)

We automatically annotate *rearing* and *grooming* behavior using JAABA (the Janelia Automatic Animal Behavior Annotator^14^, a supervised machine-learning system that performs per-frame behavior classification.

JAABA performs behavior classification using features derived from the keypoint tracking. Of the 37 landmarks, 20 were selected as the basis for single-landmark features, discarding redundant points and landmarks that were not tracked reliably. To represent pose information, we additionally defined 12 landmark pairs and 11 landmark triads that measure, for example, changes in the length of the body axis and rotation of the head relative to the body (see Supplementary Table 1 for the details of the selected landmark features). From these set of landmarks, JAABA computed 418 per-frame features: 20 landmark speeds in the arena reference frame; 200 body-centered features describing position, orientation, and velocity in the body reference frame; 132 pairwise features (distance, relative angle, velocity, area swept, and their derivatives); and 66 triad features (enclosed area, side lengths, angles, and their derivatives). Each per-frame feature was then summarized over time by JAABA’s windowed features (mean, minimum, maximum, standard deviation, harmonic content, change, difference, and z-score relative to neighboring windows).

We generated training labels manually in the JAABA interface by scrolling through the video and marking bouts of the target behavior, and bouts explicitly labeled as *not* the target behavior. For the rearing classifier we labeled 2,471 bouts (1,729 rearing, 742 not-rearing) totaling 281,821 frames (39.1 min) across 13 sessions; mean bout duration was 1.12 s for rearing and 0.56 s for not-rearing. For the grooming classifier we labeled 1,327 bouts (294 grooming, 1,033 not-grooming) totaling 301,970 frames (41.9 min) across 16 sessions; mean bout duration was 4.48 s for grooming and 1.16 s for not-grooming. Labeling proceeded iteratively: after each round of labeling, a classifier was trained, its predictions were inspected on held-out video, and additional labels were added in frames where the classifier made mistakes or was uncertain.

From the derived features and labeled frame, JAABA trains a classifier based on a boosted decision-stump algorithm ^14^. Training was done using the default training parameters (100 boosting iterations, 2,500 samples and 30 feature bins per iteration). The continuous classifier score was converted to a binary prediction using hysteresis (high 0.25, low −0.25) thresholding on the normalized score, followed by removal of bouts shorter than 20 frames (167 ms) for rearing and 60 frames (0.5 s) for grooming.

Similar to keypoint tracking, the classifier performance was assessed by ten-fold cross-validation over labeled bouts. The resulting cross-validation errors can be found in Supplementary Table 2.

### Engaged bout calculation

Engaged bouts were constructed from four event types per session: rearing bouts, tile checks, grooming, and release triggers. Rearing bouts were defined as continuous periods in which the animal stayed in one location in the arena and received one or more sound cues while rearing, as determined by the JAABA classifier. Tile checks were defined as times when the animal was inside the walls of a cricket release tile *not* at the target location, and remained there >0.5s, such that at least one sound cue was played from the correct location. This is sufficient information to indicate to the mouse that the trap door would not open at the currently visited release tile. Grooming bouts were defined as continuous periods in which the animal stayed in one location and included frames labeled as grooming by the JAABA classifier. While we created the classifier for grooming originally, it includes eating behaviors, as noted in the JAABA section of the methods. Therefore, both grooming and eating ended our engaged bouts. An engaged bout is a temporally continuous chain of rear and tile check events, ended by grooming, release triggers, or a timeout (>60s without a tile visit and >20s without a rearing event). Rears within 21-60s do not influence the current bout.

### Declaration of generative AI and AI-assisted technologies

During the preparation of this work, the authors used Claude (Sonnet 5) for generating alternative phrasings of sentences and shortening text. We also used Claude for code assistance, in particular for video generation. After using this tool, the authors reviewed and edited the content as needed and take full responsibility for the content of the publication.

## SUPPLEMENTARY FIGURES

**Supplementary Figure 1:**
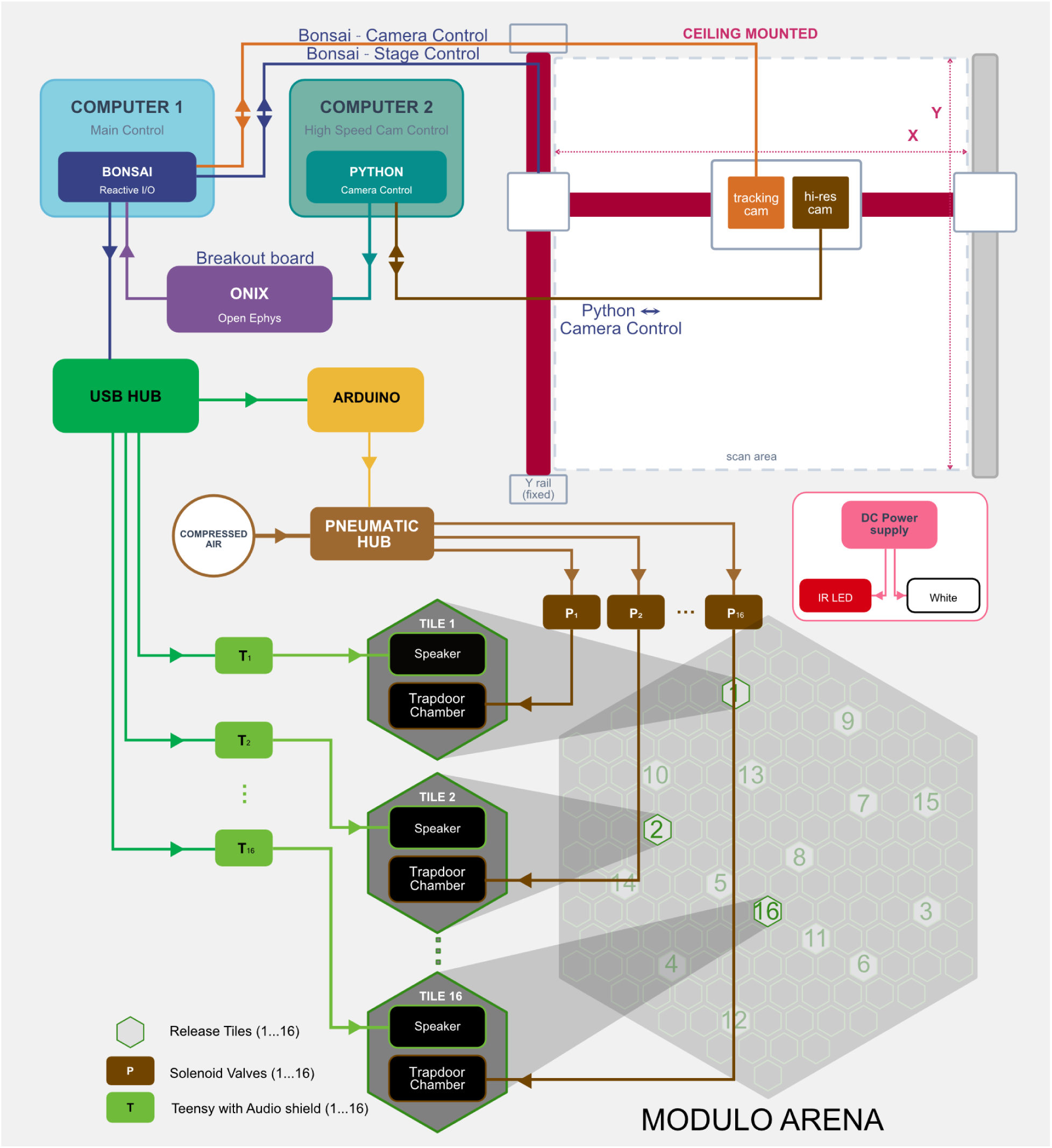
Hardware and software architecture of the Modulo system. Two computers are used in each experiment: Computer 1 (Blue, Main Control) runs a Bonsai program (Dark blue, reactive I/O), and Computer 2 (Teal: High Speed Cam Control) runs a Python application for camera control and acquires frames through a FrameGrabber at 120fps. The tracking camera feeds into Computer 1 running Bonsai, which runs a SLEAP model within the Bonsai workflow to perform live pose tracking of the animal. The resulting position estimate drives the xy stage control and tracking-camera acquisition (Dark blue: Bonsai–Stage Control and Bonsai–Camera Control paths). Bonsai drives the ceiling-mounted two-axis gantry so that the tracking camera follows the animal in real time across the arena. Bonsai on Computer 1 also controls trap door pneumatic actuation and audio, routed through a USB hub. The hub connects to (i) an Arduino that drives a pneumatic hub, which routes compressed air through solenoid valves assigned to each tile (P_1_…P_16_) to open the trap-door chambers one at a time based on the animal’s location and speed, and (ii) a bank of Teensy audio shields (T_1_…T_16_) that drive each speaker individually, allowing animal-controlled delivery of auditory cues. Each release tile therefore contains an independently addressable speaker and a pneumatically actuated trap-door chamber. Computer 2 controls the high-speed camera acquisition. An ONIX/Open Ephys acquisition system provides data synchronization across both computers. Arena illumination is provided by both infrared (IR) and white light, controlled by an independent DC power supply.

**Supplementary figure 2:**
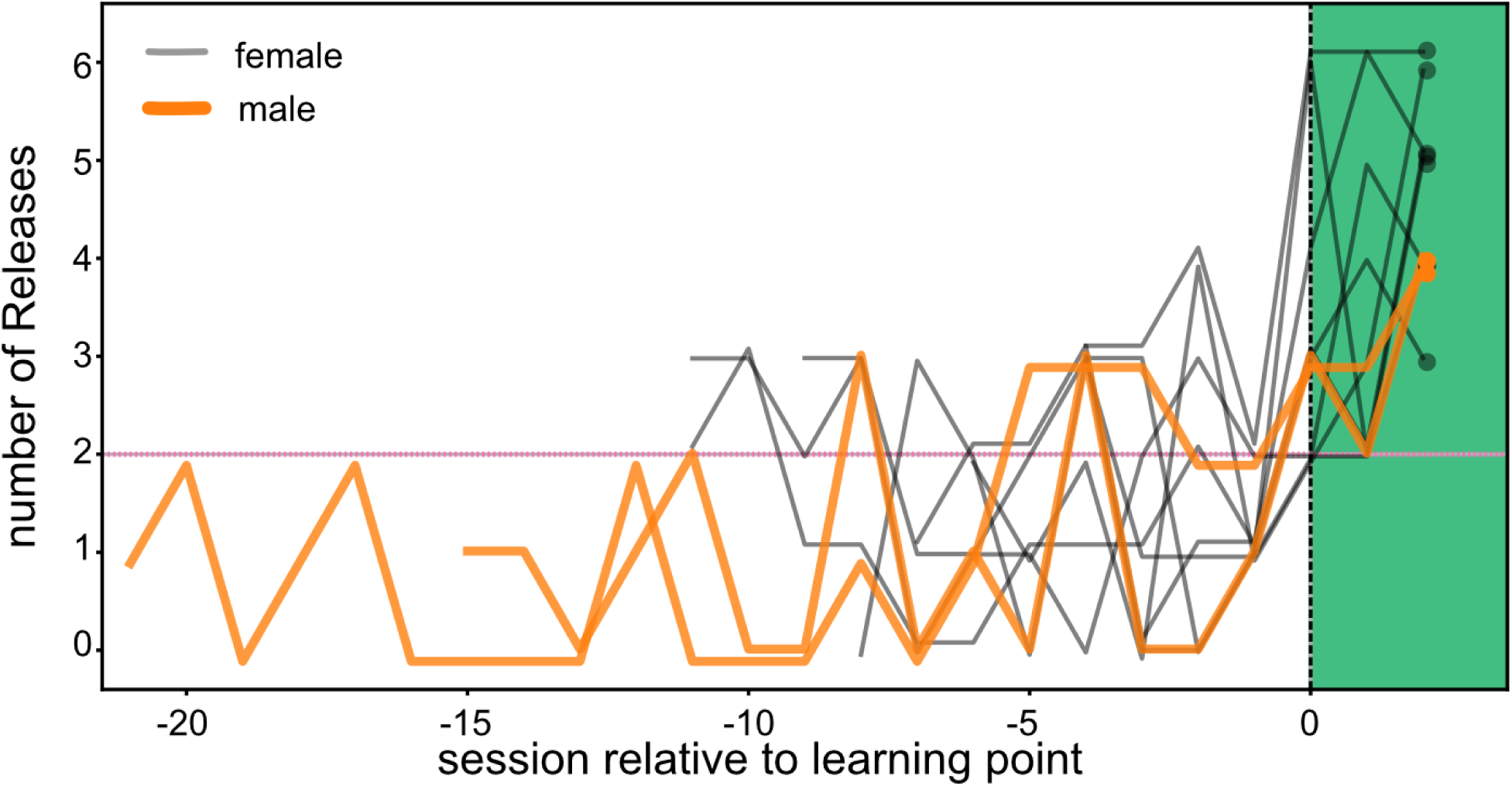
Male mice take longer to become efficient. Female Mice learn the task in 5–12 sessions (median = 8.5), data from Figure 1G. Male mice (n = 2) are plotted in orange colored lines over the female mice (n=8) in grey.

**Supplementary figure 3:**
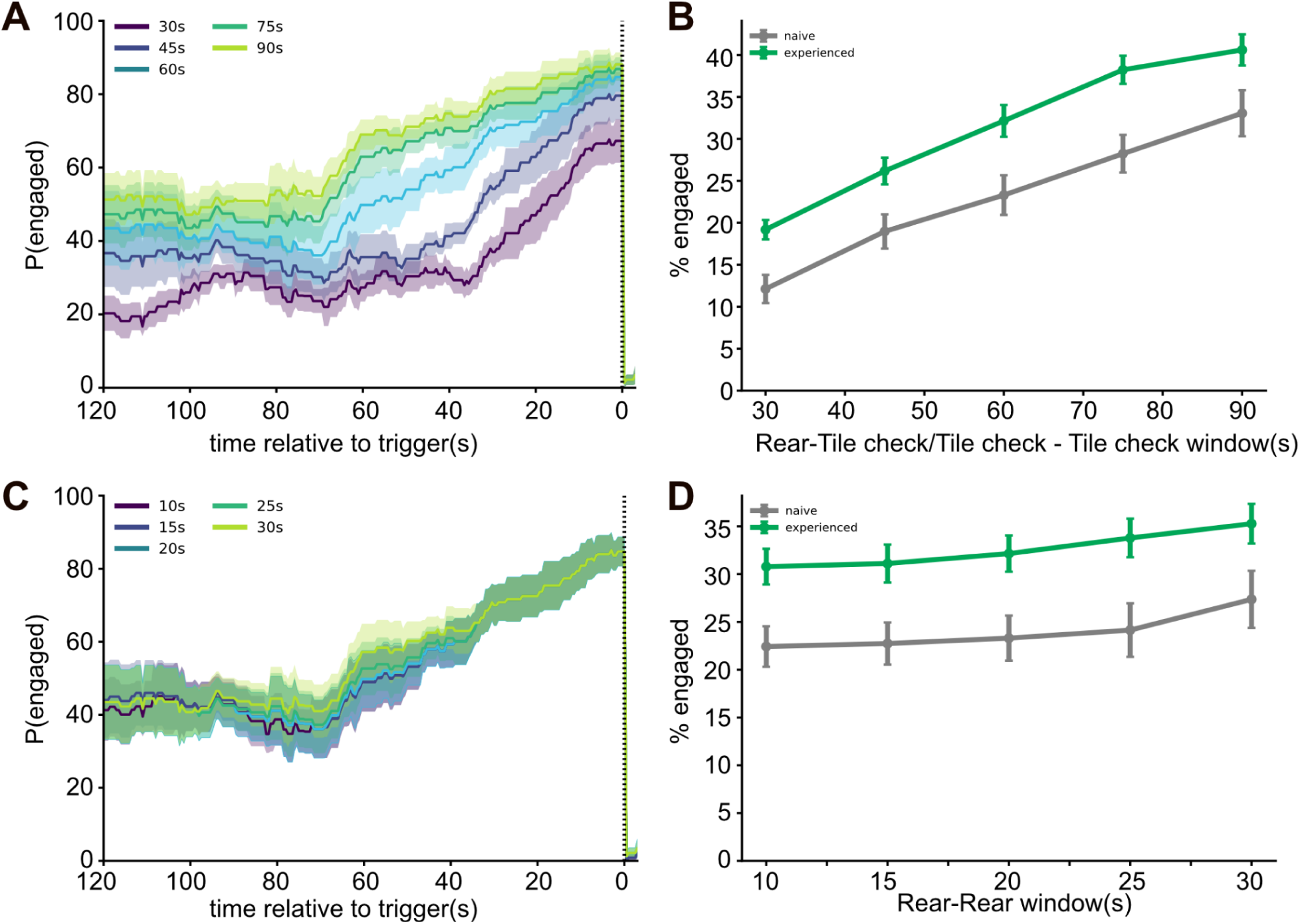
The engaged-bout structure is robust to the choice of parameters: Sensitivity of engaged-bout measures to the two temporal windows used in bout construction: the tile-check window (R-TC; the maximum interval within which a subsequent tile check is added to a bout) and the rear-rear window (R-R; the maximum interval within which a subsequent rear is added to a bout). (A) Peri-catch probability of engaged bouts as a function of time relative to the release trigger in experienced animals, plotted separately for R-TC windows of 30, 45, 60, 75, and 90 s (colours; R-R fixed at 20 s). Lines are the mean across animals and shaded bands the SEM. (B) Percentage of session time spent in engaged bouts as a function of the R-TC window, compared between naive (grey) and experienced (green) animals. Points are the median across animals (per-animal values are session medians) and error bars the SEM. (C) Peri-catch probability of engaged bouts as a function of time relative to the release trigger in experienced animals, plotted separately for, for R-R windows of 10, 15, 20, 25, and 30 s (tile-check window fixed at 60 s). (D) Percentage of session time spent in engaged bouts as a function of the R-R window, compared between naive (grey) and experienced (green) animals. Across the full range of both windows, experienced animals spent a greater fraction of time in engaged bouts than naive animals, confirming that the separation between groups does not depend on the specific window values chosen.

**Supplementary Figure 4:**
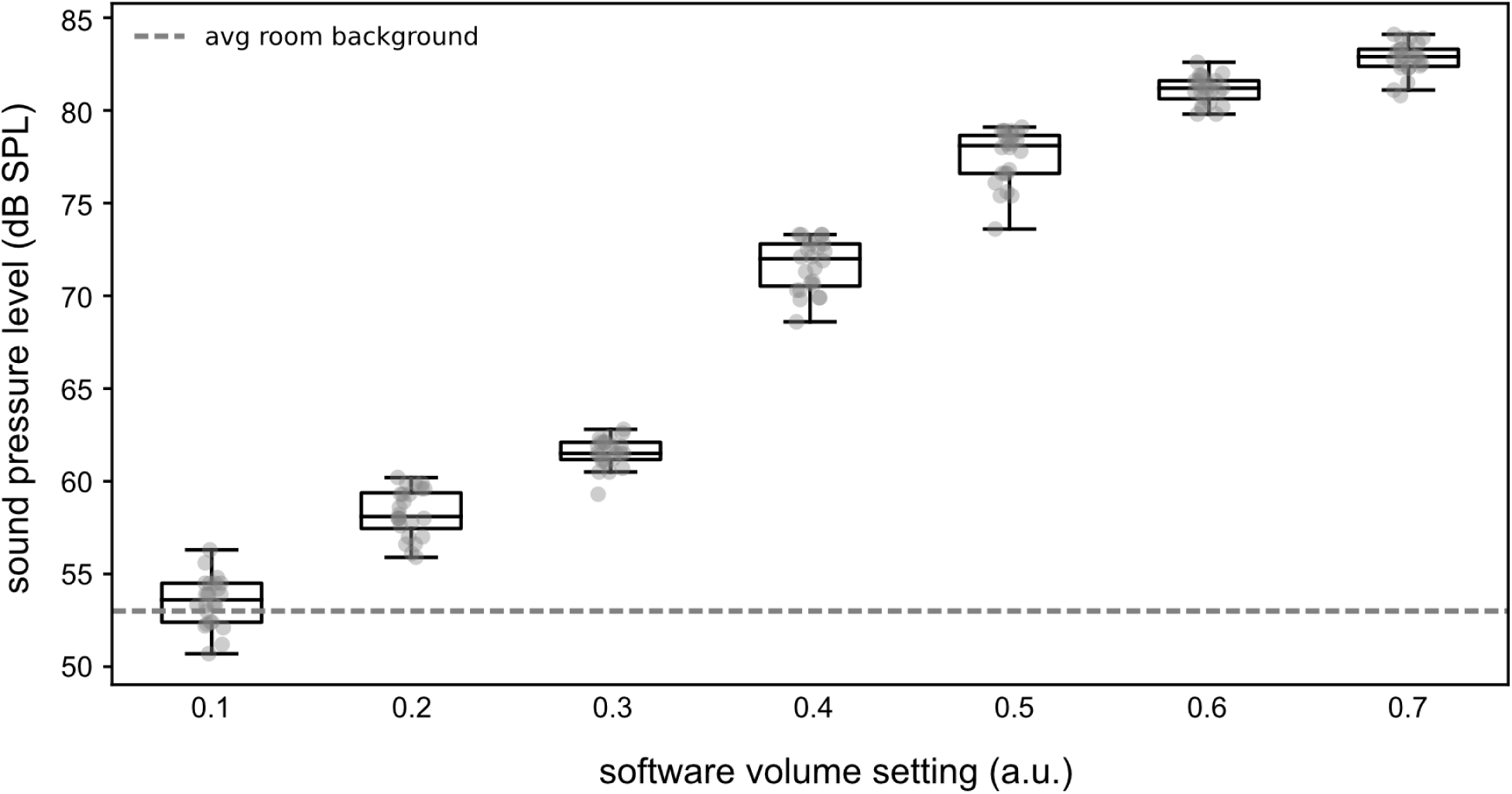
Sound pressure values (dB SPL) correlating with the software volume settings used in our experiments. Dotted line indicates room noise level.

**Supplementary figure 5:**
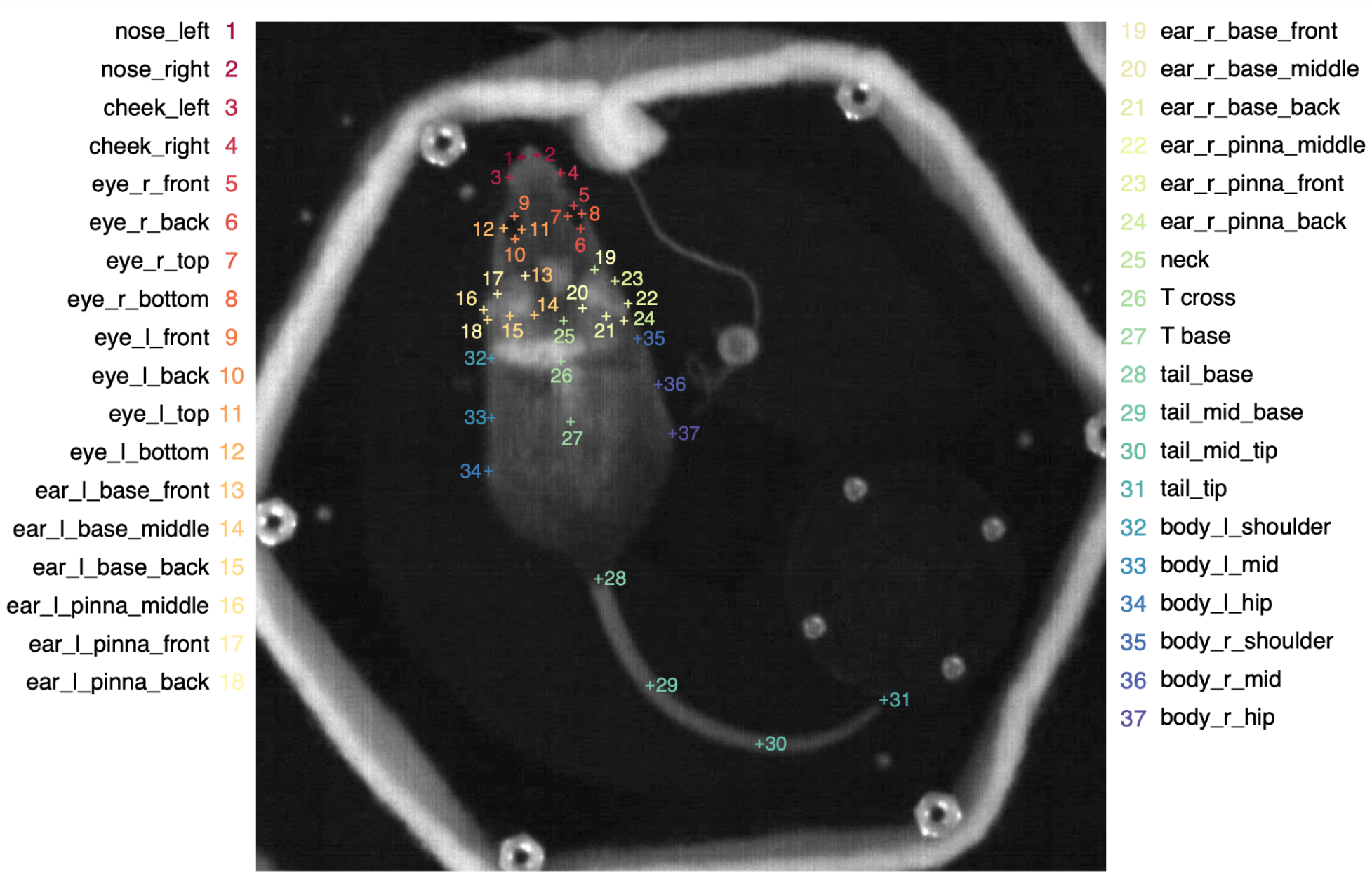
Landmarks used for keypoint tracking. We labeled a total of 37 landmarks per frame, which include the nose, eyes, ears, back, tail, and body of the animal.

**Supplementary Figure 6:**
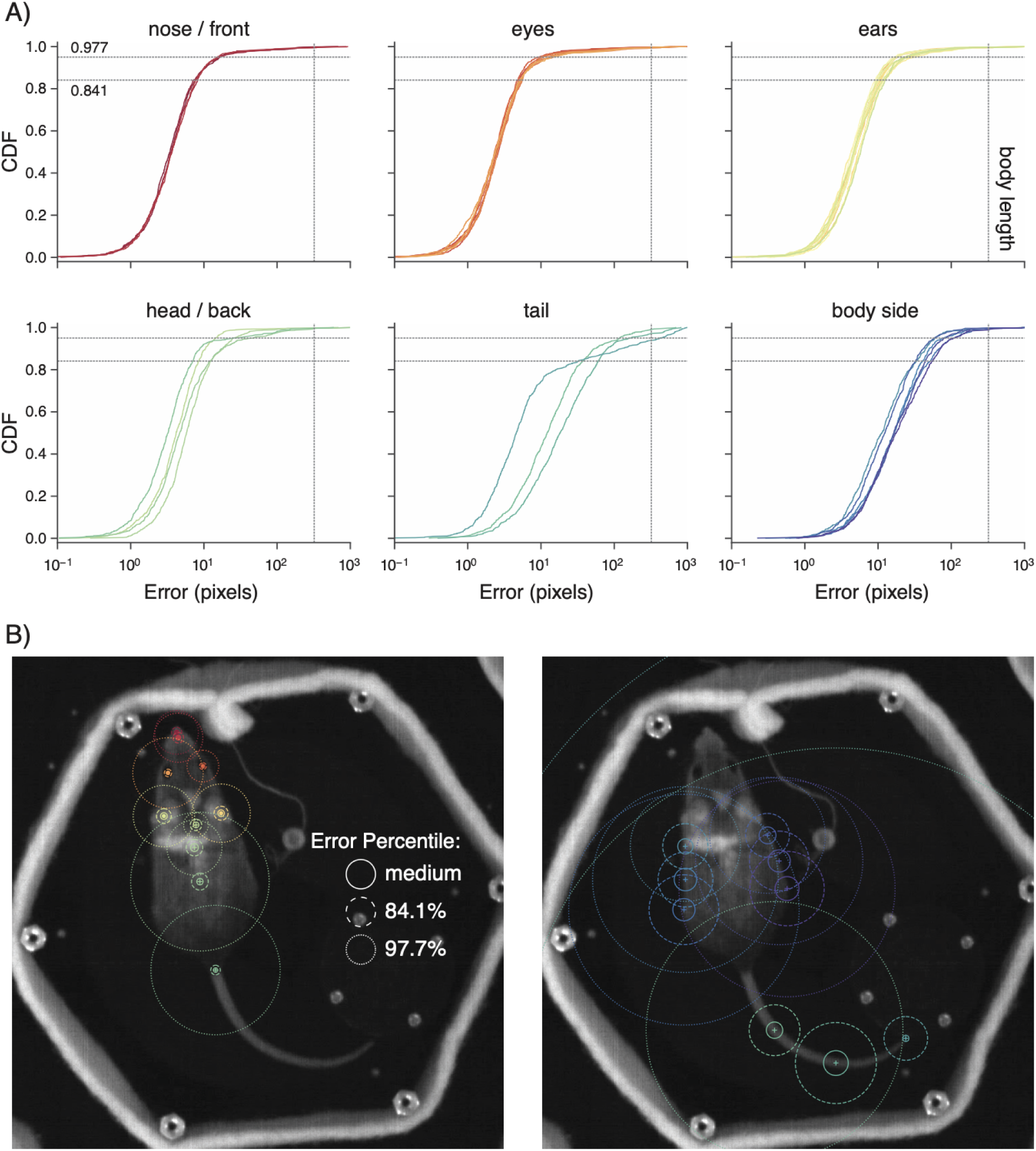
Cross-validated keypoint tracking error. (A) Cumulative distribution function (CDF) of tracking error, in pixels, for each keypoint group. The body length (vertical line) was defined as the median pixel distance between the nose and tail-base landmark. (B) Tracking error visualized on an example frame. Solid, dashed, and dotted circles denote the median, 84.1st, and 97.7th percentiles of the error distribution. Left: high-quality keypoints. Right: lower-quality keypoints used only as relational landmark pairs in subsequent analyses, not as single landmarks.

**Supplementary Table 1:** Landmarks, landmark pairs, and triads used to construct JAABA features. Related to. Figure 3.

| Feature Type | Grouping | Indices |
| --- | --- | --- |
| **Single Landmark** | Nose, Cheek | 1, 2, 3, 4 |
|  | Eyes | 5, 6, 9, 10 |
|  | Ears | 13, 15, 17, 19, 21, 23 |
|  | Body Center | 25, 26, 27, 28 |
|  | Shoulders | 32, 35 |
| **Landmark Pair** | Body Center | 25-26, 25-27, 25-28, 26-27, 26-28, 27-28 |
|  | Nose to Center | 1-25, 1-26, 1-27, 1-28 |
|  | Head | 6-10, 17-23 |
| **Landmark Triad** | Head Angle | 3-4-26, 3-4-27, 6-10-26, 6-10-27, 15-21-27, 17-23-26, 17-23-27 |
|  | Body Bend | 26-32-34, 26-35-37 |
|  | Shoulder Span | 27-32-35, 28-32-35 |

**Supplementary Table 2:** JAABA cross-validation performance. The first two columns are the frame-level confusion matrices for the rearing (top) and grooming (bottom) classifiers. Balanced accuracy is the average of the true positive and true negative rates. Precision is the fraction of predicted positive frames that are correct; Recall is the fraction of labeled positive frames that are correctly predicted as positive.

| Label / Predicted | Rearing | Not Rearing | Balanced Accuracy | Precision | Recall |
| --- | --- | --- | --- | --- | --- |
| Rearing | 216,294<br>(93.4%) | 15,335<br>(6.6%) | 92.9% | 98.3% | 93.4% |
| Not Rearing | 3,774<br>(7.5%) | 46,418<br>(92.5%) |  |  |  |

**Supplementary Table 2: JAABA cross-validation performance.**
| Label / Predicted | Grooming | Not Grooming | Balanced Accuracy | Precision | Recall |
| --- | --- | --- | --- | --- | --- |
| Grooming | 130,320<br>(82.4%) | 27,813<br>(17.6%) | 89.0% | 95.3% | 82.4% |
| Not Grooming | 6,481<br>(4.5%) | 137,356<br>(95.5%) |  |  |  |

